# Chemistry and Structure of Birch Bark Support Passive Radiative Cooling

**DOI:** 10.64898/2026.08.27.747190

**Authors:** Raffaele Perrotta, Mingna Liao, Pengli Li, Monica Ek, Florian Schott, Stephen Hall, Magnus P. Jonsson, Anna Lintunen, Mikael S. Hedenqvist, Ravi Shanker, Anna J. Svagan

## Abstract

White-barked birches extend to the northern limit of tree growth, and their bark is known to reduce solar damage during winter and early-spring by limiting solar heating and the incidence of harmful freeze-thaw events. The physical basis for this protection, however, has remained unclear. Here, we show that extracted betulin, the dominant triterpenoid responsible for the bark’s whiteness, and Himalayan birch bark, both exhibit passive radiative cooling. Under low solar irradiance, bark and betulin-pellets reach temperatures below that of a shaded reference, and pellets cool more than bark. The cooling arises from high solar reflectance, which suppresses solar heating, and substantial mid-infrared emission, which drives radiative heat loss toward outer space. These findings help explain how bark-whiteness may contribute to protecting birch trees from solar-induced thermal stress.

---

Evolution has produced organisms with exceptional passive thermoregulatory strategies for survival in extreme climates. One such strategy is passive radiative cooling (*1, 2*), in which an organism’s outer surface, such as its skin, combines two complementary processes: reflecting a large portion of sunlight in the visible and near-infrared wavelength range, which minimizes solar heating, and efficient radiative heat dissipation. In the second process, outer space acts as a natural cold sink (∼3 K). Thermal radiation escapes through the Earth’s atmosphere *via* the mid-infrared atmospheric transparency window (∼8-14 μm), allowing heat to radiate into space and enabling the organism’s temperature to fall below the surrounding air temperature, even under direct sunlight (*1–3*). This cooling strategy has been directly documented in a limited number of biological systems, most clearly in several insects inhabiting extreme thermal environments, such as the golden longicorn beetle *Neocerambyx gigas*, and in structurally white or silvery organisms, including Saharan silver ants and the butterfly *Curetis acuta* Moore (*1, 4, 5*). In plants, leaves commonly have high thermal-infrared emissivity; often around 0.95-0.98 in the atmospheric transparency window, permitting radiative heat loss alongside latent cooling by water evaporation and sensible heat exchange by convection (*6–9*).

Birch trees (genus *Betula*) are native to the Northern Hemisphere, with natural distributions across Europe, northern Asia, and North America, where many species are adapted to cold and subarctic climates (*10*). Birches typically have thin, smooth, white or varicolored bark that peels horizontally into thin sheets, particularly in young trees (*11*). The characteristic white color of several birch species is associated with abundant bark triterpenoids, with betulin identified as the dominant compound (*12–15*). Betulin concentrations are consistently higher in bark than in other plant organs, particularly in the phellem layer, reaching up to 45 wt.% of dry outer bark (*16*). The bark’s high betulin content has been suggested to serve ecological and bioactive functions, including plant defense (*17*), and its concentration varies with climate (*18, 19*), geography (*18*), and season. Guo et al. (*18*) reported higher betulin accumulation under harsher climatic conditions (colder, drier, high-latitude sites), suggesting an adaptive response to environmental stressors, although the underlying physiological mechanism was not directly tested. Separately, bark color has been shown to affect cambium temperature under natural winter conditions, with white bark stabilizing cambium temperatures, limiting solar-induced tissue damage and freeze-thaw events (*20*). However, the thermal emission properties of birch bark in the atmospheric transparency window, which are critical for passive radiative cooling, have not previously been reported.

Through a series of experiments and simulations, we demonstrate that Himalayan birch bark and a pure betulin-based material exhibit passive radiative cooling. Birch bark had an average broadband solar reflectance of 0.71 (across the 0.3-2.5 μm wavelength range) and a mid-infrared emissivity of 0.8 within the atmospheric window, producing a temperature decrease of Δ*T* = 4°C to 5°C relative to the shaded air temperature inside the testing enclosure at 0.24 sun (solar irradiance, 1 sun = 1000 W m^-2^). To isolate the role of betulin, we extracted, purified, and pressed it into porous pellets to evaluate its performance as a standalone biogenic radiative-cooling material. Concentrating purified betulin into porous pellets raised the average broadband solar reflectance to 0.89 (within 0.3-2.5 μm), while maintaining similar average mid-infrared emissivity (0.8), producing a larger temperature drop of Δ*T* = 5°C to 6°C at 0.3 sun. By directly probing the optical and thermal properties of both birch bark and purified betulin, we propose a materials-level mechanism linking composition and structure to radiative thermoregulation in birch bark.

## Morphology of birch bark and a porous betulin sample

Thin sheets of outer bark (89 ± 4 μm, mean ± s.d., Fig. 1, A , B and C) were peeled from Himalayan birch (*Betula utilis var. jacquemontii*). Examination of the bark’s interior revealed a hierarchical porous architecture, with densely packed phellem cells whose lumina were filled by triterpenoid-rich nanosheets (Fig. 1, D and E). The described morphology is consistent with previously reported triterpenoid domains in birch bark cells (*21*). These nanosheets assembled into a disordered, space-filling network lacking preferential orientation, producing a heterogeneous microstructure with micrometer-scale porosity (Fig. 1, D and E). We reason that this porosity, together with the refractive index contrast between solid and void phases (Δ*n* ≈ 0.50), would promote multiple scattering of visible light, thereby enhancing broadband reflectance and contributing to the bark’s characteristic whiteness (*22, 23*). Betulin itself is optically non-absorbing across the visible spectrum (fig. S1); its absorbance is instead confined to the ultraviolet, arising from the alkene groups within its structure (*24*).

**Fig. 1.**
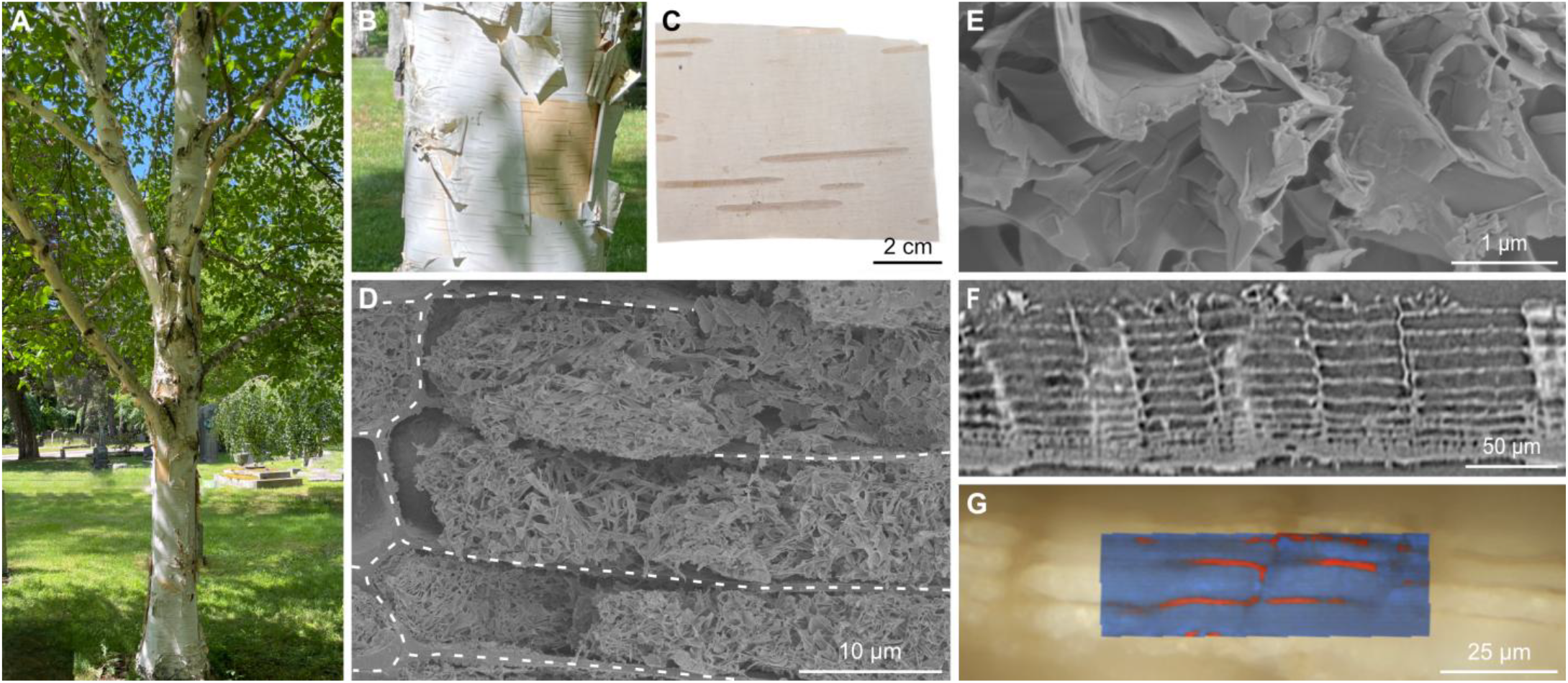
Morphology of Himalayan birch bark. (**A**) The Himalayan birch tree and magnified photograph of its stem (**B**); source of the bark in (**C**). (**D**) Scanning electron microscopy (SEM) image of densely packed phellem cells filled with triterpenoid-rich nanoscale sheets; dotted white lines mark cell walls. (**E**) High-magnification SEM of betulin nanosheets within a single phellem cell. (**F**) Longitudinal cross-section from a reconstructed µCT volume showing the phellem cells. (**G**) Color-coded Raman masks overlaid on bright-field optical image of phellem cells: red, cell wall; blue, betulin nano-sheets, identified by Raman signal.

Micro-computed tomography (µCT) revealed that most phellem cell lumina were filled, with only a minor fraction remaining empty (Fig. 1F, fig. S2). To identify this filling, we turned to confocal Raman imaging, which confirmed the content as betulin and resolved its spatial distribution within the bark. Bright-field imaging distinguished cell walls from a bright internal filling, and true component analysis (TCA) of the corresponding Raman spectra separated the characteristic signatures of the cell wall and the betulin phase, allowing us to construct a color-coded heatmap of their distribution overlaid on the optical image (Fig. 1G). Whereas cell wall spectra contained overlapping contributions from cellulose, hemicellulose, and lignin, betulin-rich domains were identified by their characteristic vibrational features at 1644, 1440, and 1195 cm^-1^ (*25*). The lignin-associated band between 1711 and 1602 cm^-1^, in particular, provided a clear point of differentiation between structural and triterpenoid components (*26*). These spectral assignments are consistent both with previously reported values and with our own Raman measurements of purified betulin (fig. S3) (*25*).

To mimic the interior architecture of birch bark cells and isolate the role of its dominant molecular component, we extracted and purified betulin, then reassembled it into free-standing porous pellets (fig. S4; Fig. 2A). Compression of betulin nanocrystal powders produced a heterogeneous structure with micrometer-scale pores (Fig. 2B), which we characterized by µCT to estimate the pellet’s pore size distribution (Fig. 2C, fig.S5). µCT-analysis identified pore diameters ranging from 2 to 20 μm, with a mean of 4.8 ± 2.1 μm (Fig. 2D); a broader distribution than that observed in native bark, with apparent pore sizes roughly around 1 μm (Fig. 1D).

**Fig. 2.**
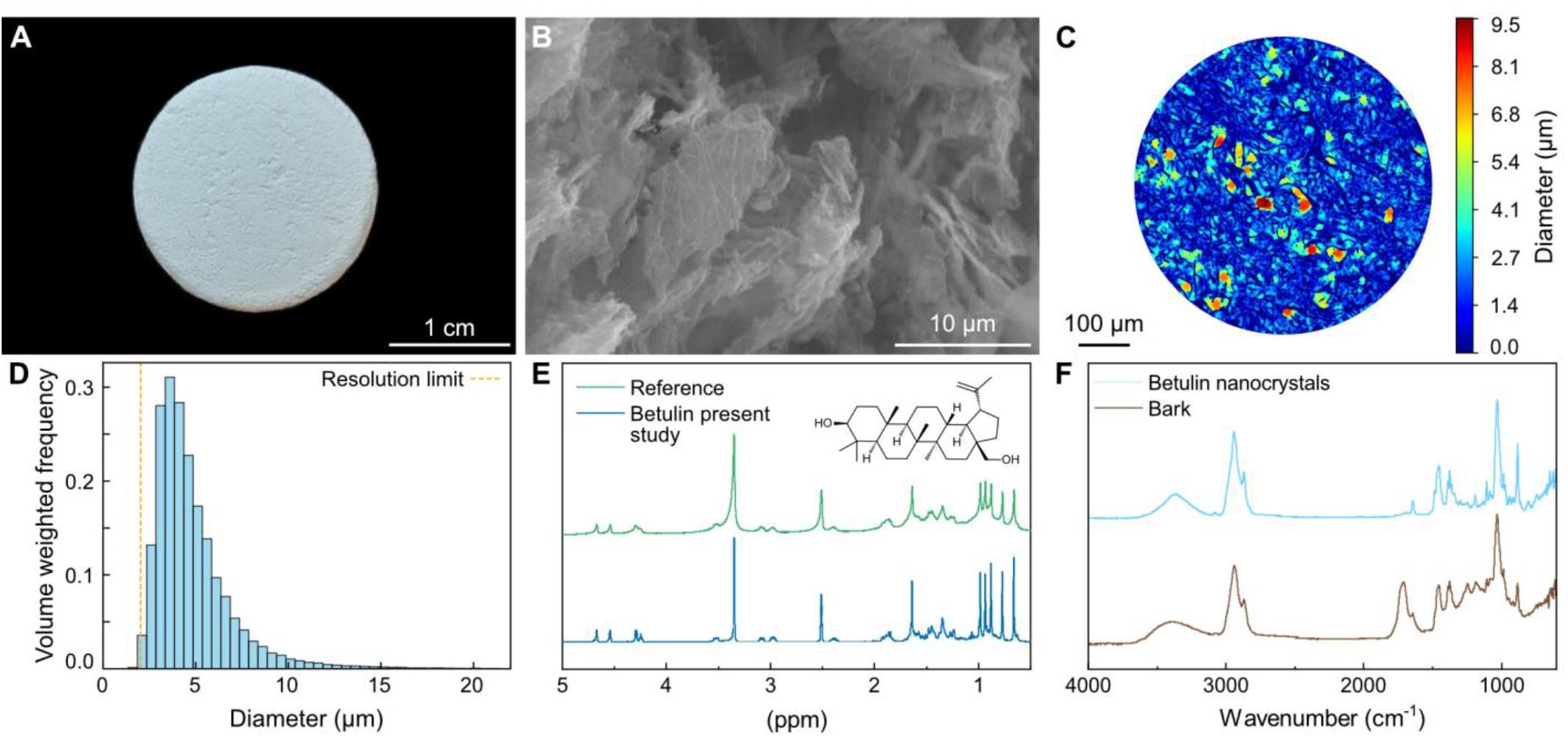
Chemistry and morphology of the betulin pellet and bark. (**A**) Photograph of a pure betulin pellet. (**B**) SEM micrograph of the pellet’s cross-section. (**C**) Typical cross-section of a µCT volumetric image of the pellet after pore-thickness quantification; the color scale indicates the diameter of the largest sphere fitting within each pore, from dark blue (solid, no pore space) to red (largest pores). (**D**) Spherical pore size distribution extracted from the µCT volumetric analysis. (**E**) ^1^H NMR spectra of recrystallized betulin and a high purity betulin reference. (**F**) FT-IR spectra of betulin used for pellet fabrication and of Himalayan birch bark.

We next verified the chemical purity of the extracted betulin using complementary spectroscopic and thermal analyses (Fig. 2, E and F; fig. S6). The proton nuclear magnetic resonance (^1^H NMR) spectrum of the recrystallized fraction matched that of a high-purity reference, displaying the characteristic proton resonances of betulin (Fig. 2E; assignments in table S1). Fourier transform infrared (FT-IR) spectra corroborated these results, with all samples exhibiting the expected vibrational signatures of betulin (Fig. 2F; assignments in table S2), confirming effective removal of impurities. For comparison, we also analyzed native birch bark. Because the mid-infrared emissive properties are governed by molecular vibrations, we note that betulin contains O-H, C-O, C-H and C=C bonds with absorption bands (table S2) falling within the atmospheric transparency window (8-14 μm; ≈ 714 to 1250 cm^-1^). The birch bark spectrum contained the same vibrational features, albeit with broader bands and additional contributions from cellulose, hemicellulose, and lignin. These results establish that both purified betulin and native bark possess the intrinsic mid-infrared vibrational modes needed to support radiative heat dissipation.

## Optical properties of betulin pellets and native birch bark

The betulin pellet scattered incident light strongly, producing high broadband reflectance across the ultraviolet, visible and near-infrared (UV-Vis-NIR) range, with values close to 100 % across much of the solar spectrum and only a slight decrease in the ultraviolet (blue curve, Fig. 3A). Averaged over the full 0.3 to 2.5 μm range, reflectance reached 0.89. This is substantially higher than the values for biological systems such as silver ants or the longicorn beetle, whose reflectance remains below 0.8 in the UV-Vis region (*1, 4, 5*). Even so, such biological reflectance still sufficed to suppress solar heat absorption. In our pellet, the suppression was even more pronounced. Its thickness and internal porosity (Fig. 2B) together generated sufficient scattering interfaces to eliminate optical transmittance (Fig. 3B), leaving absorptance below ∼5 % across most of the solar spectrum (blue curve, Fig. 3C) and indicating that the betulin pellet will effectively limit photothermal heating. The residual absorptance observed in the ultraviolet and at longer near-infrared wavelengths fell in regions of comparatively low solar irradiance and will thus contribute negligibly to heat gain. Together, the combination of high reflectance and low broadband absorptance across the 0.3-2.5 µm range provides conditions favorable for radiative cooling.

**Fig. 3.**
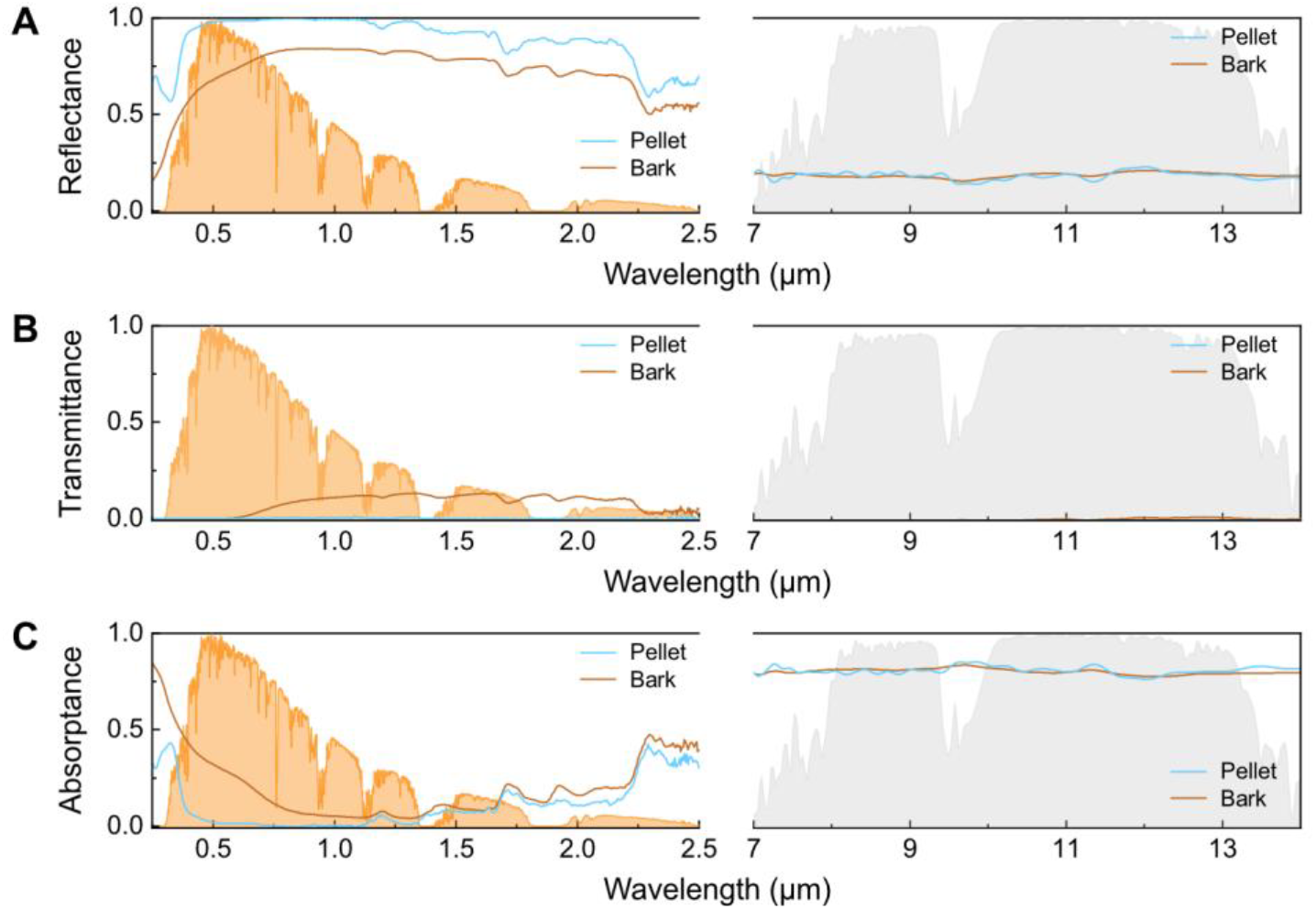
Solar and mid-infrared spectra of birch bark and the betulin pellet. (**A**) Reflectance, (**B**) transmittance, and (**C**) absorptance of Himalayan birch bark and the betulin pellet across 0.25 to 14 µm. Orange shading, solar spectral irradiance; grey shading, atmospheric transparency window.

Optical properties of the bark were measured only on the white bark regions (fig. S7), avoiding darker stripes that contain lenticels, which are static pores that pierce the bark and allow gas exchange between the stem and the atmosphere (*27*). Lenticels contain less betulin than the surrounding tissues (*28*). The bark showed a comparatively lower average reflectance of 0.71 (0.3 to 2.5 μm), which we attribute to its reduced thickness (∼89 μm) and to the presence of lignocellulosic components, particularly lignin, expected to contribute to absorption in the UV-Vis regions (*29*). Despite this lower value, the bark still provided substantial reflectance, most likely owing to its triterpenoid-rich microstructure.

The bark and pellet exhibited similar mid-infrared emissivity within the atmospheric window (0.8 for both; Fig. 3C), consistent with their shared molecular composition (Fig. 2F).

## Passive radiative cooling of bark and betulin pellet are comparable under low solar irradiance

To evaluate passive radiative cooling performance, we conducted daytime outdoor measurements of birch bark (fig. S8) and the betulin pellet (Fig. 2A) under clear-sky conditions. Each material was placed directly on a thermocouple inside a reflective enclosure, minimizing parasitic absorption from the surroundings while preserving direct radiative exchange with the sky. An infrared-transparent polyethylene film served as a wind barrier, suppressing convective losses without impeding thermal radiation (Fig. 4A). Solar irradiance was monitored in parallel (Fig. 4, C and D). Time-resolved temperature profiles of each material were compared against a reference thermocouple shaded from direct solar radiation inside the enclosure (Fig. 4, E and F; Fig. 4A). Grey shading marks intervals when a shutter covered the enclosure (Fig. 4, E to H), blocking both incident solar radiation and radiative emission and thereby equilibrating sample and enclosure temperatures. Removing the shutter exposed the sample to the sky, initiating radiative cooling and generating a temperature difference relative to the reference, Δ*T* = *T*_sample_ – *T*_ambient_ (Fig. 4, G and H). Under a solar irradiance of ∼0.3 sun (∼300 W m^-2^), the betulin pellet remained several degrees cooler than the reference throughout the measurement, reaching a maximum temperature difference of Δ*T* ≈ 5 °C to 6 °C. Under slightly lower solar irradiance (∼0.24 sun), the bark reached a temperature drop of Δ*T* ≈ 4 °C to 5 °C. This radiative cooling arises from the combination of high solar reflectance, which suppresses solar heating, and substantial mid-infrared thermal emission to the sky through the atmospheric window. Because bark and pellet share similar mid-infrared emissivity (Fig. 3C), differences in cooling performance instead primarily reflect differences in solar reflectance and thermal load. The sustained temperature reduction over time confirmed that the balance of radiative and non-radiative heat fluxes remained favorable for net heat loss under the present solar irradiance conditions, a conclusion further supported by cooling power density calculations (fig. S9A and S10A).

**Fig. 4.**
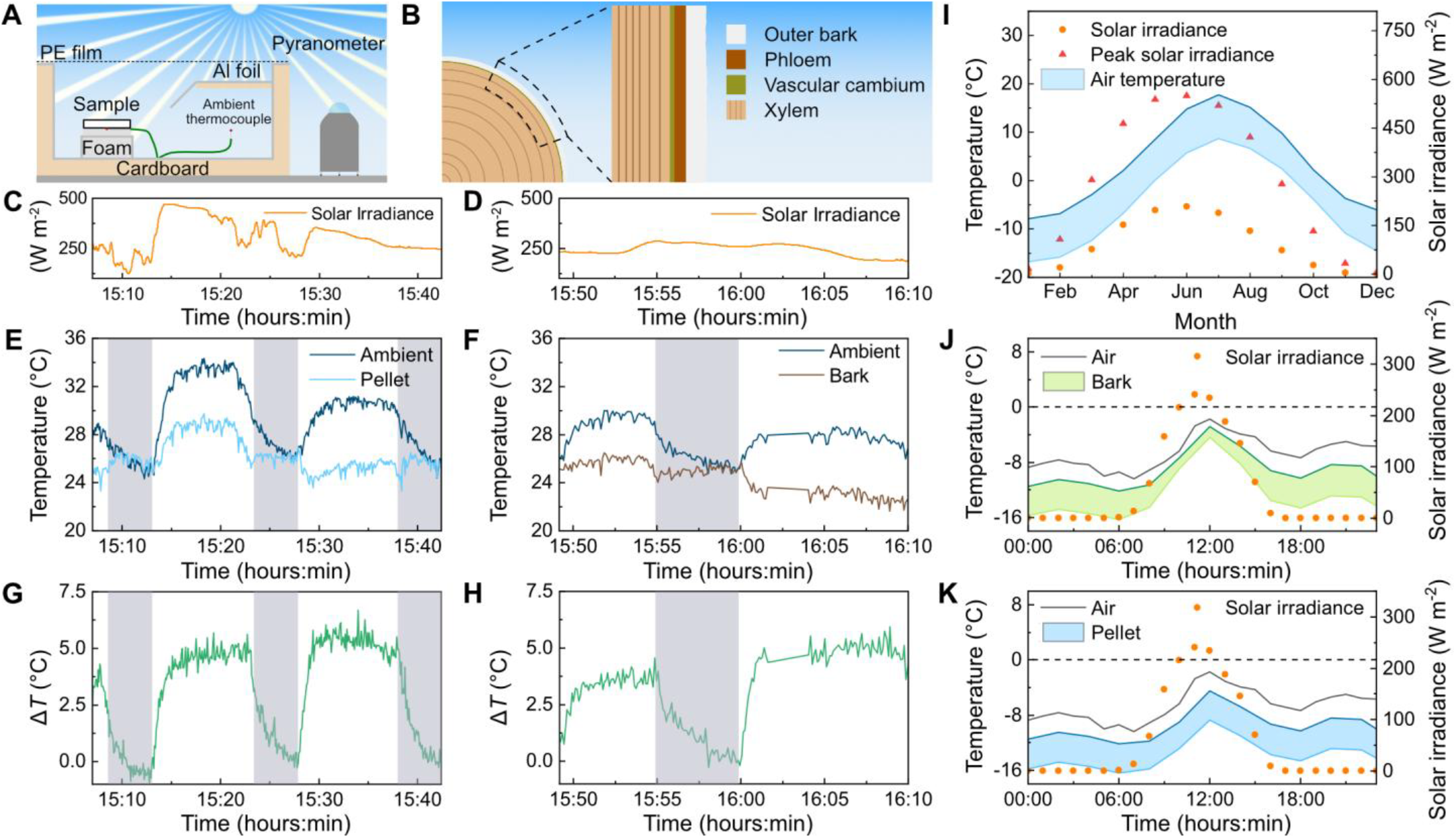
Outdoor passive radiative cooling performance of the betulin pellet and Himalayan birch bark. (**A**) Schematic of the outdoor measurement setup, with reflective enclosure and IR-transparent wind barrier. (**B**) Simplified tree stem cross-section; vascular cambium enlarged for clarity. (**C** and **D**) Real-time solar irradiance during outdoor measurements of the betulin pellet and Himalayan birch bark, respectively. (**E** and **F**) Temperature evolution of the betulin pellet and Himalayan birch bark, respectively, and the shaded ambient reference. Grey shading, shutter closed. (**G** and **H**) Resulting daytime radiative cooling of the betulin pellet and the Himalayan birch bark, respectively, showing the temperature difference relative to the shaded reference (Δ*T*). (I) Monthly average minimum and maximum air temperature (blue area) and average- and peak-solar irradiance in Kiruna (67°51’N), northern Sweden (1983-2026). Simulated temperature of (**J**) Himalayan birch bark (green area) and (**K**) the betulin pellet (blue area) during a representative sub-zero day in Kiruna (28 February 2026). Black line, air temperature; orange circles, solar irradiance, reported as a function of time.

Sunscald is an injury arising from large, rapid temperature fluctuations in the bark that kill the cambium: the layer of cells responsible for secondary stem growth, located just beneath the outer bark (Fig. 4B). It typically develops as elongated lesions or cankers on the sun-facing side of tree stems following repeated winter and early-spring solar exposure (*30, 31*). At sub-freezing air temperatures, sunscald is thought to occur when the solar radiation locally heats the bark, raising cambium temperature well above the surrounding sub-zero air. Then, when the sunlight is suddenly obstructed (e.g. shading by cloud), the temporarily thawed cambium refreezes, causing cellular death (*32–36*). The damage eventually causes the bark to become loose and fall off (*32*) and, where it extends to living cambium tissue, can act as an entry point for pathogens that indirectly cause tree death (*37*). Sunscald is most frequently observed in deciduous tree species with thin, dark bark (*30, 32, 37*).

In the boreal region, high solar radiation combined with sub-zero air temperatures primarily coincides with the tree’s leafless periods, e.g., in early spring, and Kiruna (67°51’N), a city in northern Sweden, was selected to illustrate this seasonal combination of high solar irradiance and sub-zero temperatures (Fig. 4I). To test whether the present bark limits a temperature rise and thus mitigates sunscald risk, we simulated the bark surface temperature under conditions representative of a day with potentially high sunscald risk in Kiruna; the solar irradiance peaked at ∼250 W m^-2^ with air temperature remaining below 0 °C (Fig. 4J). Under these conditions, our simulations showed that the bark was not heated above surrounding air temperatures (green area, Fig. 4J), which is important for minimizing the risk of sunscald. To identify when the bark’s cooling effect broke down, we performed additional cooling power density calculations at an artificiallyelevated irradiance of 0.5 sun under sub-zero conditions (fig. S10B); here, solar heating overwhelmed the bark’s capacity to remain sub-ambient. However, in Kiruna, an irradiance of 0.5 sun typically coincides with air temperature above 0 °C (Fig. 4I), when sunscald is not a concern.

The Himalayan birch bark sample used in this study was collected in Stockholm, so its predicted cooling power may therefore underestimate that of bark harvested from trees native to the boreal region. As a previous study showed, betulin concentration in birch bark increases at higher latitudes, where harsher climate conditions prevail (*18*). We therefore used the betulin pellet as an upper-bound reference for birch barks with enhanced reflectance. The simulated pellet temperature likewise remained below air temperatures in Kiruna, results presented in Fig. 4K (blue area). Additionally, and contrary to the bark sample, the pellet was simulated to achieve sub-ambient cooling at 0.5 sun under sub-zero conditions (fig. S9B).

## Betulin and its role in the bark adaptation to cold climate conditions

The light-colored bark of boreal deciduous trees is hypothesized to be a cold-climate adaptation that reduces the trees’ susceptibility to sunscald. Winter solar heating of bark and the resulting cambial temperature fluctuations are widely recognized as the primary contributors to sunscald injury (*20, 30–32*). Karels and Boonstra (2003) investigated the functional significance of the white bark of paper birch (*Betula papyrifera*) during winter through a series of field (Ontario, Canada) and laboratory experiments (*20*). Birch tree stems were painted with two colors (white and brown) and cambium temperatures were monitored and compared with untreated stems. On a sunny winter day with subfreezing conditions, brown-painted bark raised cambium temperatures 17.8 °C above the surrounding air temperature and above 0 °C for about three hours. Far less temperature variation (a couple of degrees above the air temperature) was observed for white-painted and untreated bark, with cambium remaining well below 0 °C. Following shading, darkened bark also cooled more rapidly. Karels and Boonstra (2003) repeatedly found that the cambium under white bark experienced smaller temperature fluctuations than cambium under darker colored bark during normal winter conditions. Long-term field observations (2 years) further revealed that brown-painted trees developed markedly more sunscald injury than trees with natural or white-painted bark, supporting the hypothesized protective role of light-colored bark. Additionally, frequent freezing and thawing of a tree stem can increase the risk of winter embolism formation and thereby lower the efficiency of water transport in a tree in the following growing season (*38*). Karels and Boonstra (2003) concluded that white bark limits winter solar heating of the cambium, lowering freeze-thaw stress and sunscald risk, and proposed this as an adaptation contributing to why white-barked species, among deciduous trees, are the only ones to reach the northern limit of tree growth in North America (*20*).

In the present study, we show that birch bark can cool passively through radiative heat loss under specific conditions. By combining structural, chemical, optical and thermal analyses, we found that the tested white bark samples couple high broadband solar reflectance with substantial thermal radiation toward outer space. The latter represents an additional and previously overlooked mechanism. We propose that this heat-loss pathway contributes to thermal regulation during clear winter days, when solar radiation becomes sufficient to warm the bark while air temperatures remain below freezing. Under such clear-sky conditions, radiative cooling helps keep bark temperatures closer to ambient, thereby lowering the likelihood of thawing followed by rapid refreezing of the stem and the associated risk of cambial tissue damage and winter embolism.

A previous study of *Betula platyphylla* across Northeast China found that bark betulin and total triterpenoid contents were higher at high-latitude sites, a pattern attributed in part to colder, drier geo-climatic conditions (*18*). Our experiments with extracted betulin, further suggest that betulin is an important contributor to the thermoregulatory function. Concentrating purified betulin into a highly scattering porous material considerably boosted solar reflectance while maintaining mid-infrared emissivity comparable to native bark, yielding stronger cooling performance. To conclude, our observations suggest that a high betulin content, characteristic of white birch bark, not only contributes to its visible whiteness, but also provides the material properties necessary to regulate temperature *via* passive radiative cooling.

## Materials and methods

### Materials

Ethanol (96 %), dimethyl sulfoxide-d6 (DMSO, 99.96 atom % D) and heptane (>99.5 %) were purchased from VWR Chemicals, Sigma-Aldrich and Fisher Chemicals respectively. Betulin (SB) was purchased from Sigma-Aldrich (≥ 98 %). Bark was obtained from a *Betula utilis* (Himalayan birch), grown in Stockholm, ca. 30 years old.

### Betulin extraction and purification

The bark of a *Betula pendula* was cut from a fallen tree located in the Norra Djurgården forest, in Stockholm, Sweden. The outer bark (OB) was manually separated from the inner and dried at 105 °C overnight. Then, a Wiley mill (3383 L10, Thomas Scientific, USA) was used to mill the OB to a 40 mesh. The ground OB was stored at ambient condition in a sealed container. Betulin was obtained from the OB by Soxhlet extraction using heptane as solvent. A 100 mL Soxhlet apparatus was used to process 12.3 g of OB using 300 mL of solvent, which was brought to a boiling with a heating mantle. The heptane extracted betulin (HB) precipitated as a yellowish powder once the extraction set-up cooled down, and it was collected through vacuum filtration. The extraction required approximately 20 h. HB was purified by recrystallization: it was dissolved in boiling ethanol under stirring, and betulin crystals (RB) formed once the solution cooled. RB was recovered through vacuum filtration.

### Preparation of betulin nanocrystals

Betulin nanocrystals (BN) were obtained from RB through a solvent exchange precipitation process, adapting a protocol developed by Zhao et al. (*39*). Recrystallized betulin was fully dissolved in boiling ethanol 96 % and then poured in deionized water at ambient temperature under stirring. The solution was slowly poured and the betulin crystals immediately formed upon mixing. The solution was freeze-dried to obtain the betulin nanocrystals.

### Preparation of betulin pellets

A 3D printed mold in polylactide, inspired by Poh et al. (*40*), was designed to produce the betulin pellets and it was printed with an X1-Carbon (Bambu Lab, China). The mold consists of three components: a cylindrical body with a 25 mm circular opening and a height of 25 mm, a bottom platform with a protruding cylinder that fits the opening in the main body, and a cylindrical pressing element equipped with two conical lateral guides to control its travel. The mold was designed to produce betulin pellets with a diameter of 25 mm and thickness of 2 mm. Before pressing, the contact surfaces were covered with a polyethylene terephthalate anti-sticking foil, after which 180 mg of BN were added and manually pressed. Larger pellets of 4 cm in diameter were produced to satisfy the minimum sample dimension for the spectrophotometers’ integrating sphere used in the optical characterization. The mold design was modified for the pellet to be kept inside a cylindrical sample holder (fig. S7).

### Scanning electron microscopy

The morphology of HB, RB and the betulin pellet was observed with a Tabletop SEM TM-1000 (Hitachi, Japan) with a 15 kV accelerating voltage, while BN and the bark were analyzed with a Field-Emission SEM S-4800 (Hitachi, Japan) at 1 kV. The betulin pellet and the bark were sputtered for 30 s with Pt/Pd with a sputter coater (208HR, Cressington, UK).

### X-ray diffraction

The diffraction spectra of HB, RB, and BN were obtained with a X’Pert Pro X-ray diffractometer (PANalytical, Netherlands) with Cu Kα radiation with a wavelength of 1.54 Å. The angular range 2θ of 5 - 60° was analyzed in continuous mode, using a 0.04° s^-1^ scan rate. The Origin 2020 software (OriginLab Corporation, USA) was used for baseline correction.

### X-ray micro-computed tomography

The X-ray micro-computed tomography of the Himalayan birch bark and of the betulin pellet were performed with a Zeiss Versa XRM 730 (Carl Zeiss X-ray Microscopy, USA) at the 4D Imaging Lab at Lund University using source voltage of 80 kV and power of 10 W. After tomographic reconstruction of the projection data, 3D image volumes with cubic voxels of widths 855 nm, for the bark, and 675 nm, for the pellet, were achieved. The tomographic images were quantified using a standardized Python image-analysis pipeline (*41*). Background inhomogeneity was removed using a top-hat filter (*42*), followed by a median filter to suppress speckle noise (*42*). A cylindrical mask was applied to exclude peripheral regions of the images that were prone to limited-angle reconstruction artifacts. Gas pores were segmented from the solid betulin crystal phase using a global Otsu threshold, and the pore-crystal interfaces were refined with a Sobel filter and ITK watershed algorithm (*42, 43*). The local pore thickness was quantified by measuring the diameter of the largest sphere that could be inscribed within the pore space and centered at each pore voxel (*44*). Finally, a Gaussian filter was applied to the resulting thickness maps to obtain a continuous pore-diameter field. The resolution limit shown in Fig.2 is defined as three times the voxel size.

### Raman microscopy

The chemical composition of the Himalayan birch bark was analyzed using an alpha300 confocal Raman microscope (Witec GmbH, Germany). Confocal imagining was performed with a Zeiss EC Epiplan-Neofluar Dic 50×/0.8 objective. The Raman spectra were acquired using a 785 nm excitation laser with a power of 30 mW and 0.5 s integration time. The Raman signal was directed to a spectrometer (300 groves mm^-1^, UTHS300, Witec GmbH, Germany) equipped with a charge-coupled device (CCD) camera. The set-up reached a lateral resolution of approximately 500 nm. A large area scan of 70×22 μm^2^ was performed acquiring 3 spectra µm^-1^. Prior to testing, the bark sample was cryo-fractured and taped onto a vertical support. The Project SIX software (Witec GmbH, Germany) was used for the removal of cosmic rays, the background subtraction and the smoothing of the spectral data with the Svitzky Golay method using a 10-point window and a fourth order polynomial. The True Component Analysis (TCA) tool in Project SIX was used to identify the bark components and generate the color-coded heatmap.

### Fourier-transform infrared

The Himalayan birch bark and betulin functional groups were analyzed with a Spectrum 100 (Perkin Elmer, USA) with a Universal Attenuated Total Reflectance accessory equipped with a Germanium crystal. 16 scans at a resolution of 4 cm^-1^ were performed over the 4000–600 cm^-1^ wavelength range. The software Spectrum (Perkin Elmer, USA) was used to correct the baseline.

### Proton nuclear magnetic resonance spectroscopy

An Ultrashield 400 MHz (Bruker, USA) 54 mm long hold magnet (p/n BZH/422/400/70F) equipped with a B-ACS60 autosampler was used to obtain the ^1^H NMR spectra. RB and SB were dissolved in DMSO in concentrations of 5 mg mL^-1^. Undissolved particles were removed using a Nylon 0.45 μm syringe filter. The MestReNova software (Mestrelab Research, Spain) was used for data processing and baseline correction.

### Differential scanning calorimetry

The melting and crystallization of the betulin samples were analyzed with a DSC1 (Mettler Toledo, Switzerland). The measurements were performed under a 50 mL min^-1^ nitrogen flow rate over the temperature range 30-270 °C, with a 10 °C min^-1^ heating rate. Two heating and two cooling cycles were performed for each sample. The samples were loaded in 100 µL aluminum crucibles, duplicates were tested.

### Thermogravimetric analysis

The betulin thermal decomposition was investigated with a TGA/DSC1 (Mettler Toledo, Switzerland). The samples were heated with a 10 °C min^-1^ heating rate from 30 to 500 °C under a 50 mL min^-1^ nitrogen flow rate. Duplicate samples were tested in 70 µL alumina crucibles, with an empty one providing the baseline.

### Optical properties

The spectral reflectance and transmittance of the samples were measured from the ultraviolet to the mid-infrared wavelength region using two complementary instruments. A LAMBDA 1050+ (PerkinElmer, USA) spectrophotometer was used for the 250–2500 nm range, and a Spectron 3 FTIR (PerkinElmer, USA) spectrometer covered wavelengths from 7 to 14 µm. Both setups were equipped with integrating spheres to record directional-hemispherical reflectance and transmittance under near-normal incidence. For the LAMBDA system, an integrating sphere was used with PMT and PbS detectors selected for the UV–VIS and NIR regions, respectively. The FTIR measurements employed a Mid-IR IntegratIR sphere (PIKE Technologies, USA) with an MCT detector. Reflectance values were calibrated using Spectralon® (UV–VIS range, Labsphere, USA) and the Mid-IR Diffuse Reflection Wavelength Standard (PIKE Technologies, USA) reference standards. Absorptance spectra were obtained from the relation A(λ) = 1 ™ R(λ) ™ T(λ), and spectral emissivity was taken as equal to absorptance according to Kirchhoff’s law. This allowed evaluation of both solar reflectance and mid-infrared emissive properties relevant to passive radiative cooling.

### Outdoor measurement

The setup for the radiative cooling measurement consisted of a cardboard box 28 x 20 x 5 cm (L x W x H) whose surfaces were covered with reflective aluminum foil and sealed on top by a 12 μm polyethylene foil to isolate the setup from the wind. The sample stage consisted of a foam block to minimize heat exchange with the box, covered with aluminum foil and an additional piece of foam was taped on top to prevent heat conduction from the aluminum layer to the sample. The latter was placed on the upper foam block with a K type thermocouple (Pentronic, Sweden) in the middle, taped to the foam. A free thermocouple, shielded from the direct sunlight, was used to measure the air temperature in the box, considered as the reference ambient temperature. The temperature reading of the thermocouples were recorded with an Arduino microcontroller using the QuikEval software. The solar irradiance was measured with a MS-60 pyranometer (EKO Instruments, Japan). The radiative cooling measurement was performed on an outdoor rooftop at Linköping university, campus Norrköping (58°35’23.5”N, 16°10’36.1”E) on October 11, 2025. The average temperature, wind speed and relative humidity during that afternoon were 15 °C, 6 m s^-1^, and 55 %.

### Historical data of solar irradiance and temperature in Kiruna

Solar irradiance and temperature data for the city of Kiruna (Sweden) were obtained from the Swedish Meteorological and Hydrological Institute (SMHI) database (*45*). The monthly average and monthly peak solar irradiance from January 1983 to January 2026 were calculated from the SMHI hourly solar irradiance dataset (Kiruna Sol station, 67.8408° latitude, 20.4105° longitude, 423 m above sea level). The monthly average minimum and maximum temperature within the period January 1983 to January 2026 were computed from the daily minimum and maximum temperature dataset (Kiruna Airport station, 67.8270° latitude, 20.3387° longitude, 459 m above sea level).

## Acknowledgments

We thank B. Birdsong for support in the initial development and printing of the 3D printed mold. We thank E. Tyrode for support in performing the Raman microscopy measurements. The AI model Copilot assisted the coding used for the power density calculations and for the analysis of the tomographic images.

## Author contributions

Conceptualization: RP, RS, AJS

Methodology: RP, MJ, PL, SH, RS, AJS

Investigation: RP, RS, ML, PL, SH, FS

Visualization: RP, RS

Funding acquisition: MH

Supervision: AJS, MH, ME, MJ

Writing – original draft: RP, RS, AJS

Writing – review & editing: all authors

